# Direct Feature Identification from Raman Spectra and Precise Data-driven Classification of Phytopathogens at Single Conidium-Species Level

**DOI:** 10.1101/2024.11.29.626083

**Authors:** Xinze Xu, Zhang Cheng, Wenbo Liu, Chunhua Lin, Weiguo Miao, Heyang Yuan

## Abstract

Conidia can cause outbreaks of fungal plant diseases, resulting in significant losses in crop yield and economy. Traditional diagnosis of phytopathogenic conidia based on morphology and molecular biology is time-consuming, labor-intensive, and often fails to differentiate fungal conidia. To overcome these challenges, a new classification approach was developed in this study by integrating Raman spectroscopy with data-driven modeling. Eight fungal species were selected and characterized using Raman spectra. Three characteristic Raman wavenumbers at 1003 to 1005 cm^-1^, 1153 to 1157 cm^-1^, and 1515 to 1522 cm^-1^ shared a consistent pattern across species and could be attributed to carotenoids. Clustering of the Raman spectra using principal component analysis (PCA) showed substantial overlap, indicating inaccurate classification of conidia. Three data-driven models, support vector machines (SVMs), decision trees (DTs), and eXtreme Gradient Boosting Forest (XGBoost) were trained with three categories of features (number of peaks, maximum peak, and curve roughness) identified within eight characteristic wavenumber ranges, The optimal SVM, DT, and XGBoost determined by hyperparameter tuning achieved prediction precision of 0.88, 0.88, and 0.96, respectively. PCA-XGBoost trained by feeding principal components of PCA to XGBoost achieved prediction precision of 0.94, suggesting that features extracted from the raw datasets outperformed those extracted with PCA in terms of data-driven classification. This study has demonstrated the great potential of Raman spectroscopy combined with data-driven modeling for classification of phytopathogenic conidia.

## 1. Introduction

Large-scale plant pandemics could reduce global yields of major crops by up to 15%, resulting in economic losses of hundreds of billions of dollars [1]. Approximately 8,000 species of fungi and oomycetes, accounting for 70-80% of phytopathogens, have been reported as the main culprits of severe plant pandemics [2–5]. For example, conidia are a type of fungal spores that have been shown to be the primary cause of widespread plant diseases. They are capable of asexual reproduction [6], which can significantly affect phenology and increase the incidence of infection. Rapid and accurate identification of phytopathogenic fungi such as conidia is highly desired for protection of agricultural products.

Traditional methods to diagnose phytopathogenic fungal diseases are based on susceptibility symptoms, sclerotia or mycelium, spore morphology, and sporulation structures [7]. These methods require well-trained personnel with expertise in fungal taxonomy, plant pathology, and specialized operation skills [8,9], making the diagnosis costly, time-consuming, and laborious. Moreover, the information obtained from the morphology of a single conidium is often unrepresentative at the species level, which can lead to biased diagnoses [10,11]. In recent decades, diagnostic technologies based on molecular biology, such as polymerase chain reaction (PCR), have provided more precise and sensitive alternatives for detecting and quantifying plant pathogens [12,13]. However, for early detection where spores collected from infected leaves are insufficient, molecular techniques are ineffective due to low DNA concentrations. In addition, pre-enrichment in the laboratory is hindered when obligate parasitic fungi cannot be cultured on non-host media. These limitations have motivated the development of alternative methods for accurate identification of phytopathogenic fungi, particularly conidia.

Raman spectroscopy represents a promising tool for rapid, reagent-free, non-destructive, and high-throughput identification of phytopathogenic fungi [14–16]. This technique has been applied to differentiate the phenotypic traits, physiological states, and metabolic activities of cells with similar structures, including human teardrops [17] and fungi associated with moisture-damaged building materials [18]. A combination of Raman spectroscopy and confocal Raman microscopy can capture signals in the focal region and discard background noise, enabling accurate analysis of samples at the micrometer scale (e.g., 1-3 μm) such as a single cell or spore [19,20]. For example, culture-free bacteria were detected at the variant level [21], and dermatophyte fungi were identified at the genus and species levels [22]. There is a growing interest in combining this experimental technique with advanced computational approaches to improve the identification of fungal conidia.

In recent decades, data-driven modeling has emerged as a new tool for reliable identification of phytopathogenic fungi. Unsupervised data-driven methods such as principal component analysis (PCA) can extract features through dimensionality reduction and classify samples in low-dimensional space [21–23]. Supervised data-driven methods have also shown potential in classification tasks [24]. Popular supervised methods, such as artificial neural networks (ANNs), demonstrate superior learning capabilities [25,26] but require the calibration of large amounts of parameters and can learn noise (i.e., overfitting) from small training sets [27]. In comparison, lightweight supervised methods, such as supporting vector machines (SVMs) and decision trees (DTs), can yield better predictive performance when dealing with small training sets [27,28]. For example, SVMs were built to diagnose patients with dengue infection and achieved a diagnostic accuracy of 85% based on 84 samples [29]. DTs were also used to diagnose various human diseases or classify functional cell lines [30,31]. The eXtreme Gradient Boosting Forest (XGBoost), an improved version of DTs, was developed to overcome the overfitting issues of traditional DTs [32] and has been widely applied in the fields of disease diagnosis, image recognition, and text analysis, etc [33–35]. These data-driven methods could be powerful tools to extract valuable information from Raman spectra for improved identification of phytopathogenic fungi.

In this study, data-driven methods were developed to identify phytopathogenic conidia from Raman spectra. The objectives are to (1) identify the common features in the Raman spectra of conidia from different species, (2) verify the ability of PCA to capture the Raman features of conidia, (3) optimize three supervised data-driven models, SVMs, DTs, and XGBoost using features extracted from raw Raman spectra, and (4) compare the identification accuracy of XGBoost trained with raw features and with PCA-extracted features. The last objective is examined because feature extraction can play a key role in building data-driven models when the training dataset contains complicated features such as Raman spectra [36,37]. This study is expected to contribute to the understanding of the Raman spectra of phytopathogenic conidia and facilitate the development of reliable classification methods in agricultural disease management, ultimately benefiting agricultural sustainability.

## 2. Materials and Methods

### 2.1 Sample preparation

Eight species, *Fusarium graminearum*, *Colletotrichum alatae*, *Colletotrichum tropicale*, *Colletotrichum siamense*, *Colletotrichum acutatum, Erysiphe cichoracearum*, *Erysiphe quercicola*, and *Podosphaera hibiscicola*, were selected as representatives of different phytopathogenic fungi. *F. graminearum* and the four *Colletotrichum* species were cultured on potato dextrose agar at the temperature of 28. Five millimeters of sterile water was then pipetted onto the potato dextrose agar for suspension of conidia and filtered through Miracloth cloths. *E. quercicola* was cultured on young, bronze-stage leaves of the moderately susceptible rubber tree cultivar, Reyan 7-33-97 [38]. *E. cichoracearum* and *P. hibiscicola* were cultured on mature leaves of high-yield agricultural cowpea and an *Arabidopsis pad4* mutant, respectively. The conidia of the powdery mildew fungi were directly brushed off the infected leaves and collected into 2-mL centrifuge tubes. The conidial samples were then suspended by 2 mL of sterile water and filtered through Miracloth cloths (pore size 22-25 μm, Merck KgaA, Germany) to remove mycelial fragments. All filtrates of the eight species were centrifuged at 10,000 rpm for 3 minutes. The supernatant was discarded, and the pellets were re-suspended followed by centrifugation. This process was repeated two times to remove soluble impurities, and the final pellets were diluted to 10^5^ CFU/mL using sterile water.

### 2.2 Raman spectroscopy

Twenty millimeters of conidial suspension of each species, which contained 2×10^3^ conidia, was dropped onto a single-sided polished monocrystalline silicon substrate (1 cm × 1 cm) and then blow-dried for later use. Single-sided polished monocrystalline silicon was used as the substrate to minimize spectral interference because it showed a low Raman scattering cross-section and high optical transparency [39]. This material is also more cost-effective and reusable compared to other Raman spectroscopy substrates. Accordingly, a 514-nm excitation laser was selected for its compatibility with monocrystalline silicon to enhance the quality of the Raman spectra.

Raman spectra were recorded using a micro-Raman spectrometer (inVia Reflex of Renishaw, England) equipped with a 50 mW, 514 nm laser. The 520 cm^-1^ signal of monocrystalline silicon wafers was used to calibrate the Raman shifts before measurements. The laser intensity for sample irradiation was controlled at approximately 5 mW, and the diameter of the laser spot was set to 1 μm. Raman spectra were recorded from 600 to 2000 cm^-1^. This range covers most of the spectral bands that include critical fungal characteristics. A spectral resolution of 1 cm^-1^ was achieved by a 2400 mm^-1^ grating in the spectrometer. Finally, The Raman spectra of the four *Colletotrichum* conidia were obtained using a 50× objective lens, whereas *F*. *graminearum* and the three powdery mildews were examined under a 20× lens, with each measurement being recorded over a ten-second responding time. Each species was characterized with biological triplicates, and twenty spectra were recorded at random focal spots for each replicate, resulting in 60 spectra per species. To avoid potential photodamage-induced alterations of conidial spectral profiles, no repeated measurements were carried out on the same conidium. A total of 480 Raman spectra were collected for the eight species.

### 2.3 Data pre-processing

The Raman peaks of individual conidia of the same species could be inconsistent due to environmental noise and individual differences. Meanwhile, the autofluorescence background of the samples and random variations could also interfere with spectral imaging, resulting in issues such as elevated baseline and false positive peaks. Thus, before classification analysis, several data pre-processing steps were carried out to standardize the raw spectra: 1) background subtraction (i.e., monocrystalline silicon), 2) curve smoothing (four degrees), and 3) baseline correction (≥ five degrees, Polymon) using LabSpec 5 (HORIBA Scientific, Orsay, France) [40,41]. To demonstrate the error interval and intrasample repeatability of the spectra, the standard deviations of the Raman intensity were calculated corresponding to each wavenumber across the selected range, and the error margin was shaded on both sides (upper and lower area) of the normalized (mean-centered Z-score method) average spectra.

### 2.4 Data-driven identification of conidia

#### 2.4.1 Feature selection and dataset generation

After pre-processing, the spectral data were arranged in the form of relative intensity at different Raman shifts (i.e., wavenumbers). Number of peaks, maximum peak, and curve roughness were calculated as three categories of features using R (Version 4.3.2). These features were identified within eight shift ranges (i.e., 940-980, 980-1020, 1120-1200, 1260-1340, 1400-1480, 1500-1540, 1580-1620, and 1620-1680 cm^-1^) that exhibited signal peaks and/or significant fluctuations in one or more species (Fig. 1). Peaks were automatically identified when a local maximum with relatively large two-side gradients was detected. The gradients were estimated using a finite difference approach [42]:

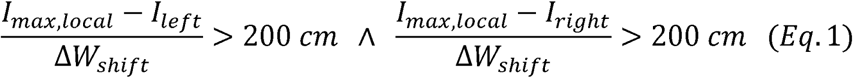

**Fig. 1.**
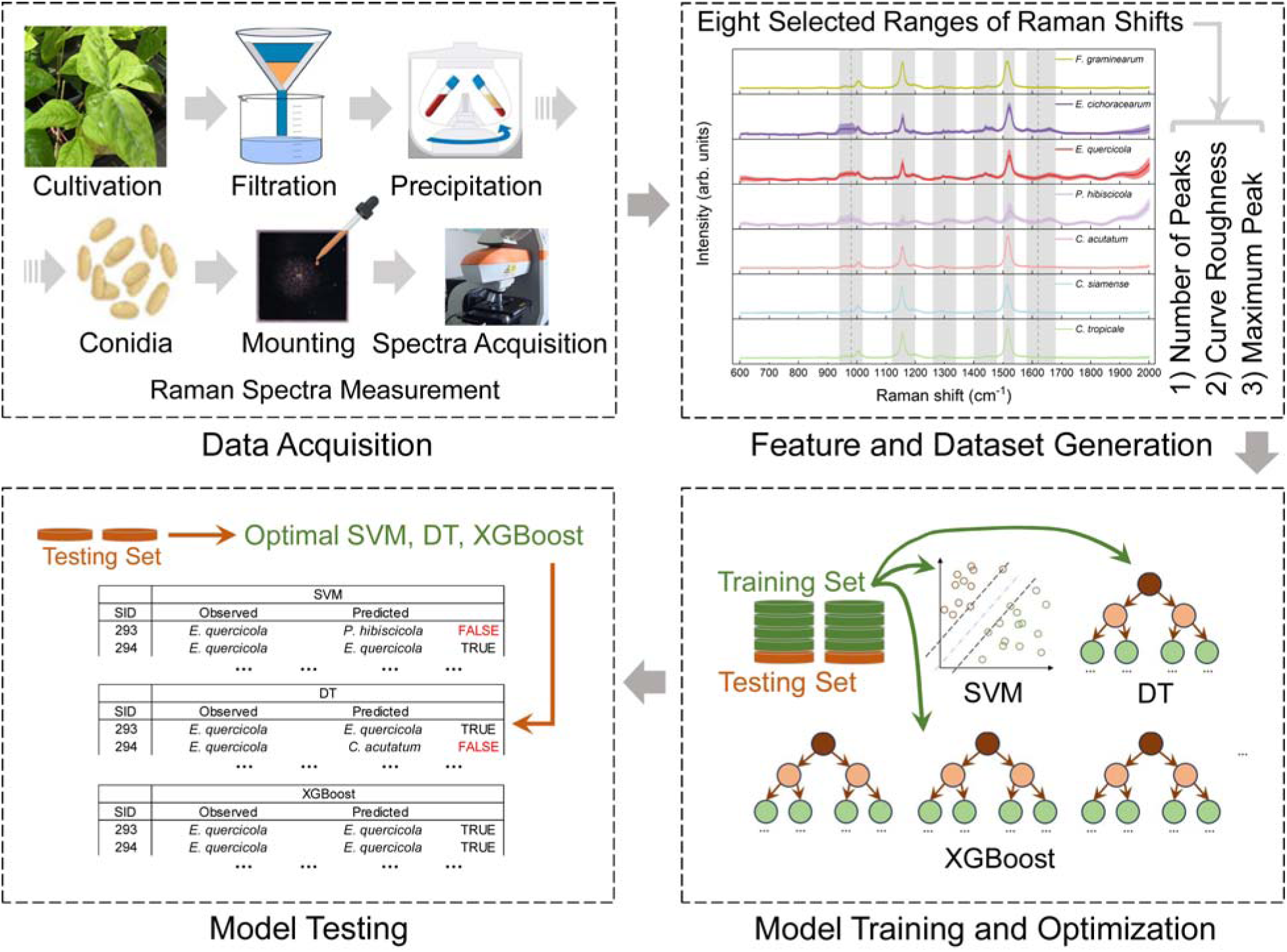
Schematic of combining Raman spectroscopy and data-driven modeling for identification of fungal conidia.

Where *1_max,local_* is the local maximum signal intensity, Δ*w_shift_* is the width of Raman shift for peak identification and set to 5 cm^-1^ in this paper, *I_left_* is the intensity of the left side with a shift of Δ*w_shift_* from the local maximum, and *1_right_* is the intensity of the right side with a shift of Δ*w_shift_* from the local maximum. The curve roughness was expressed as the standard deviation of the data within each range of the selected Raman shifts. The transformed samples were rearranged under the 24 (3×8) features, with labels indicating the corresponding phytopathogenic fungal species (Supplementary File 1).

Before training data-driven models, the values under the three categories of features (i.e., number of peaks, maximum peak, and curve roughness) were scaled to 0 – 1 [43]:

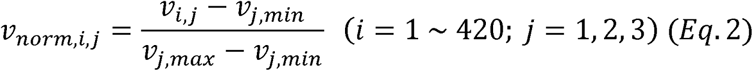

Where *v_i,j_* is the value *i* under category *j*, *v_j,max_* is the maximum value under category *j*, *v_j,min_* is the minimum value under category *j*, and *v_norm,i,j_* is the normalized data *i* under category *j*. For each species, the training and test sets consisted of 48 and 12 samples (Fig. 1), respectively.

#### 2.4.2 PCA analysis

PCA was performed to cluster the species. The distances between pre-processed spectra were calculated using the covariance matrix and visualized with two components. Clustering analysis was performed twice. The first PCA was based on the entire spectral range and the corresponding normalized Raman intensities. The second one was based on the features generated from the eight characteristic Raman shift ranges, as described above. Both PCA results served as references for the other three data-driven methods built as described below.

#### 2.4.3 Construction of data-driven models

Three R packages, e1071, rpart, and xgboost, were used to construct SVMs, DTs, and XGBoost, respectively. PCA-XGBoost was also trained with a dataset composed of principal components derived from PCA [44,45]. All model trainings were performed on the Temple High Performance Computing with a computational configuration of 80 CPU cores. The training set was first shuffled, and a six-fold cross-validation was performed to optimize the hyperparameters and verify the model robustness [46]. The hyperparameters that were tuned included the kernel function for SVM, the maximum depth for DT, and the maximum depth and learning rate for XGBoost. Unless specified, all other hyperparameters were set to defaults in the R packages. The detailed settings for training the data-driven models can be found in Table 1. The micro-average precision (MAP) was calculated to evaluate the results of either the cross-validation or the test set [47]:

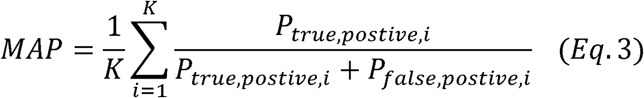

**Table 1.**
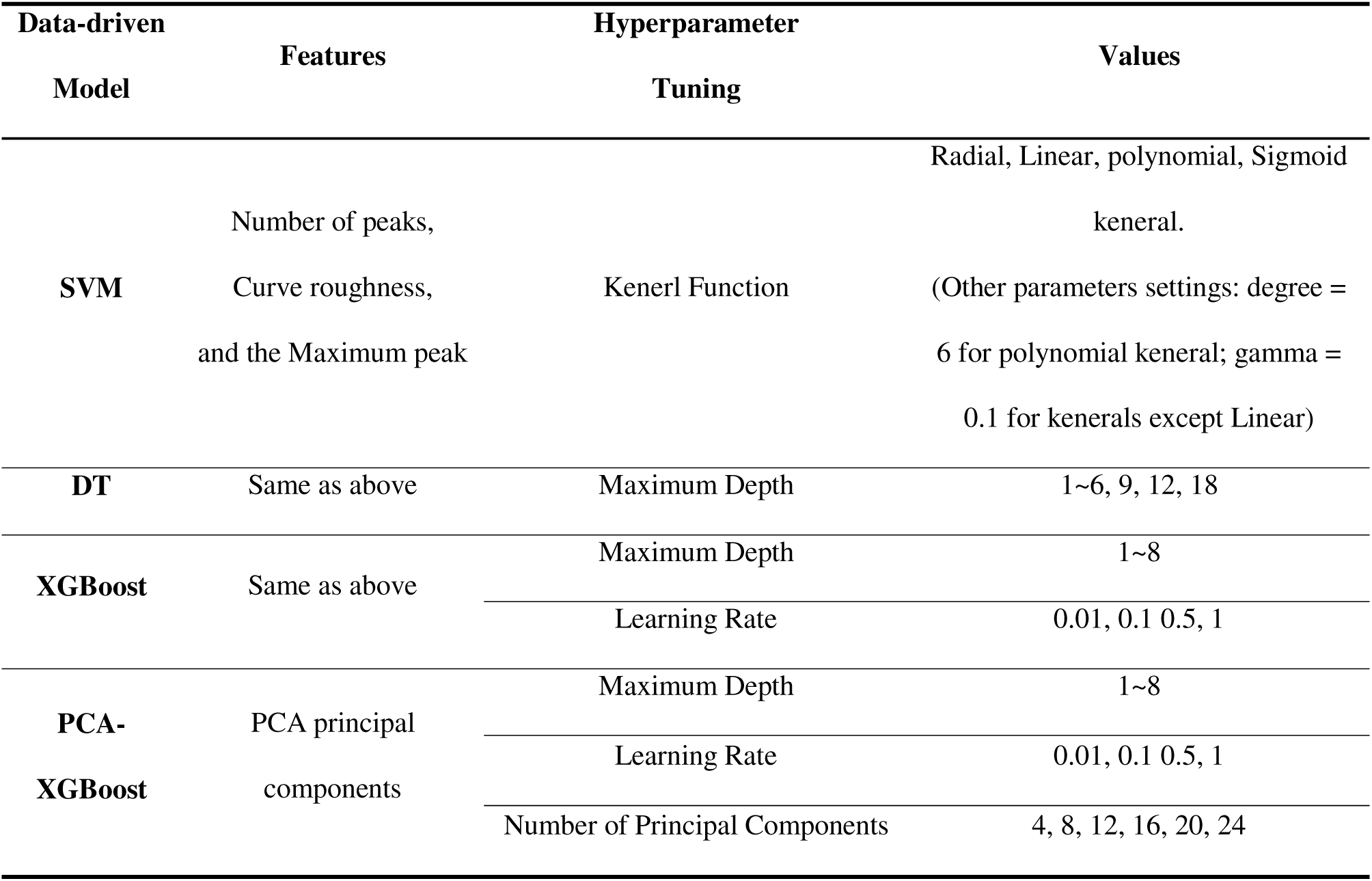
The training settings of data-driven models.

Where *p_true,postive,i_* is the successful prediction of positive samples from fungal species *i*, *p_false,postive,i_* is the false prediction of positive samples from fungal species *i*, *K* = 7 is the number of the labels (i.e., fungal species). In the context of multiclass classification, it is important to note that when a specific class is evaluated, all other classes are considered negative. Given that the same number of samples from each species were used to compose the training and testing set, the MAP equals the overall accuracy. Finally, the optimized models were tested against the test set.

## 3. Result and Discussion

### 3.1 Raman spectral characteristics

The selected fungal species can be divided into three main groups: 1) *F*. *graminearum* is a severe plant pathogen posing a significant threat to crops; 2) *C*. *tropicale*, *C*. *siamense*, *C*. *acutatum*, and *C*. *alatae* belong to the genus *Colletotrichum*, and 3) *E*. *quercicola*, *E*. *cichoracearum*, and *P*. *hibiscicola* are powdery mildews. Although the samples from different genera differed in morphology, host, pathogenicity, and living environment, their Raman spectra showed similar patterns, especially with primary peaks around 1005 cm^-1^, 1156 cm^-1^, and 1518 cm^-1^ (Fig. 2). For example, the full range spectra of *F*. *graminearum* and three of the four *Colletotrichum* species were highly similar. It should be noted that *C. alatae* produced strong Raman signals and exceeded the detection limitation despite adjustment of the laser power. The reason for the abnormal signals was not clear, and *C. alatae* was not included in the subsequent data-driven analysis. The three powdery mildews also showed spectra highly analogous to the other two groups with slight differences observed. These included a significant spectral oscillation and standard errors of the primary peaks indicated by the shaded regions (Fig. 2). The elevated trend in the spectra of the powdery mildews after 1625 cm^-1^ was possibly due to the autofluorescence background of conidia that raised the baseline of the original spectra. These subtle differences might not allow accurate, reliable, and rapid identification of powdery mildews at the species level. The results suggested a consistent composition of endogenous substances and, consequently, the difficulty of visual discrimination of different species via Raman spectra.

**Fig. 2.**
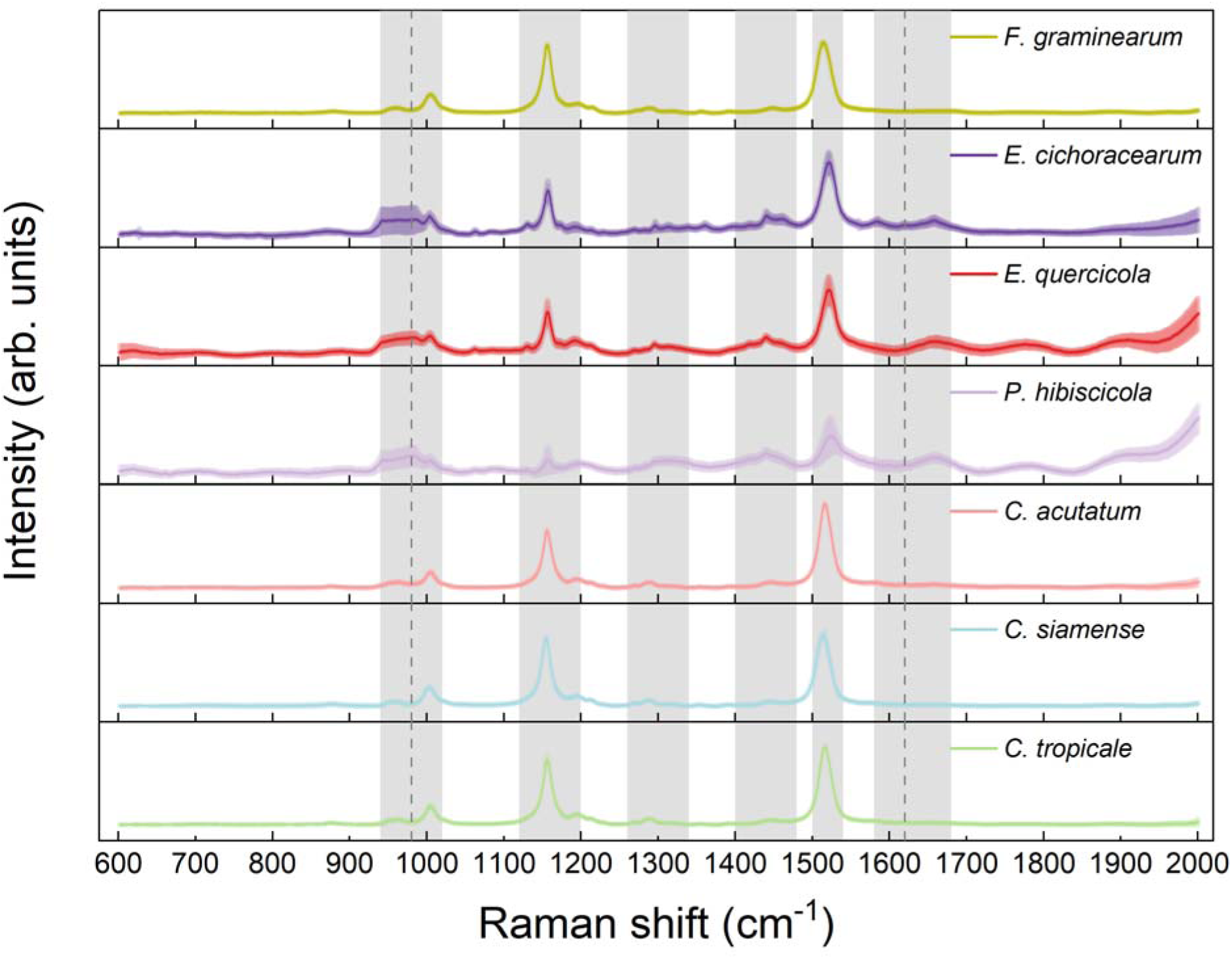
The Raman spectra of the *F*. *graminearum,* three powdery mildews (i.e., *E*. *cichoracearum*, *E*. *quercicola*, and *P*. *hibiscicola*), and four *Colletotrichum* species (i.e., *C*. *alatae*, *C*. *acutatum*, *C*. *siamense*, and *C*. *tropicale*). The spectra were obtained by averaging the 60 spectra of individual species. The resolution is 1 cm^-1^, and the shaded area represents the range of standard deviation.

The primary peaks within 1003 - 1005 cm^-1^, 1153 - 1157cm^-1^, and 1515 - 1522 cm^-1^ of all spectra could be contributed by C-H, C-H_3_, C-C bend [48,49] C-C stretching vibration[48,49]; and C=C stretching vibration [50,51], respectively. In early studies, the peak at 1003 - 1005 cm^-1^ was also considered to be related to the ring breathing mode of phenylalanine in the protein of the cytoplasm [52,53]. In comparison, our primary peaks were likely to be closer to the Raman patterns of carotenoids such as β-carotene and astaxanthin. This is because the spectra of the conidia in this study are highly similar to the Raman signatures of *Haematococcus pluvialis*, an alga rich in astaxanthin and β-carotene [51]. The varying intensities of the primary peaks might indicate the distinct contents of carotenoids among different species. Carotenoids have been widely reported in phytopathogenic fungi and are significant antioxidants that can help microbes resist oxidative stress, protect them from light-induced damage, and enhance their pathogenicity [54,55]. Significant carotenoid production could provide insights into the interaction between pathogens and hosts through quantitative analysis of the relationship between Raman intensity/width and metabolism/transcription [56,57].

Secondary peaks were identified to be those within 1190 - 1194 cm^-1^, 1290 - 1295 cm^-1^, and 1439 - 1446 cm^-1^, which could represent C-H deformation vibration of β-carotene and amide III [58], C-C bend of amide III [59], and C-H_2_ deformation vibration of unsaturated fatty acid [50,60], respectively (Table 2).

**Table 2.** The band assignments of major Raman peaks observed in conidial Raman spectra of all species.

| Range/cm<sup>-1</sup> | Band assignment |
| --- | --- |
| 960-963 | C-H<sub>3</sub> |
| 940-990 | Silicon second-order Raman scattering |
| 1002-1005 | C-H, C-H<sub>3</sub>, C-C bend |
| 1153-1158 | C-C stretching vibration |
| 1190-1194 | C-H deformation vibration of β-carotene and amide III |
| 1290-1295 | C-C bend of amide III |
| 1439-1446 | C-H<sub>2</sub> deformation vibration of unsaturated fatty acid |
| 1515-1527 | C = C stretching vibration |
| 1656-1661 | Amide I |

These secondary peaks could result from various substances. For example, the peak at 960 - 963 cm^-1^ was reported as a representative of C-H_3_ [20], while the 940 - 990 cm^-1^ was also recognized as the Raman shifts representing secondary peaks of monocrystalline silicon. Hence, the peaks measured from 930- 995 cm^-1^ were not assigned to any specific substance in this study. In conclusion, despite the presence of slightly distinct patterns in the spectra, the similarities in spectral trends and peaks continue to pose challenges for visual discrimination of the fungal conidia solely via the raw Raman spectra. The development of robust classification tools is imperative.

### 3.2 Principal component analysis

PCA was performed on the whole spectral range of a total of 420 conidial spectra of seven species (excluding *C. alatae* due to abnormal spectra). As shown in Fig. 3a, the samples could be divided into two clusters based on 95% confidence intervals, with the first and second principal components contributing 68% and 11% of the explained variance, respectively. Specifically, the first principal component could explain the peaks at 1154, 1513, and after 1858 cm^-1^ (Fig. S1), while the patterns at 979, 1151, 1508, 1525, and 1659 cm^-1^ were mainly reflected by the second principal component. The PCA results did not allow a clear identification of *F*. *graminearum*, as this species showed considerable overlap with *C*. *siamense* and *C. tropicale*. On the other hand, the three *Colletotrichum* species and three powdery mildews demonstrated genus-level differentiation. *Colletotrichum* could also be visually distinguished at the species level although the 95% confidence interval partially overlapped. For the three powdery mildews, *E. cichoracearum* and *P. hibiscicola* were separated with minor overlap, while *E. quercicola* overlapped with the other two species to an indistinguishable extent. Clustering can be improved by including more components. As shown in the PCA scree plot (Fig. 3b), the eigenvalue at the inflection point was five, and the first five principal components could account for 93% of the cumulative explained variance (Supplementary File 2). More accurate clustering could be achieved in high-dimensional spaces but could not be visualized.

**Fig. 3.**
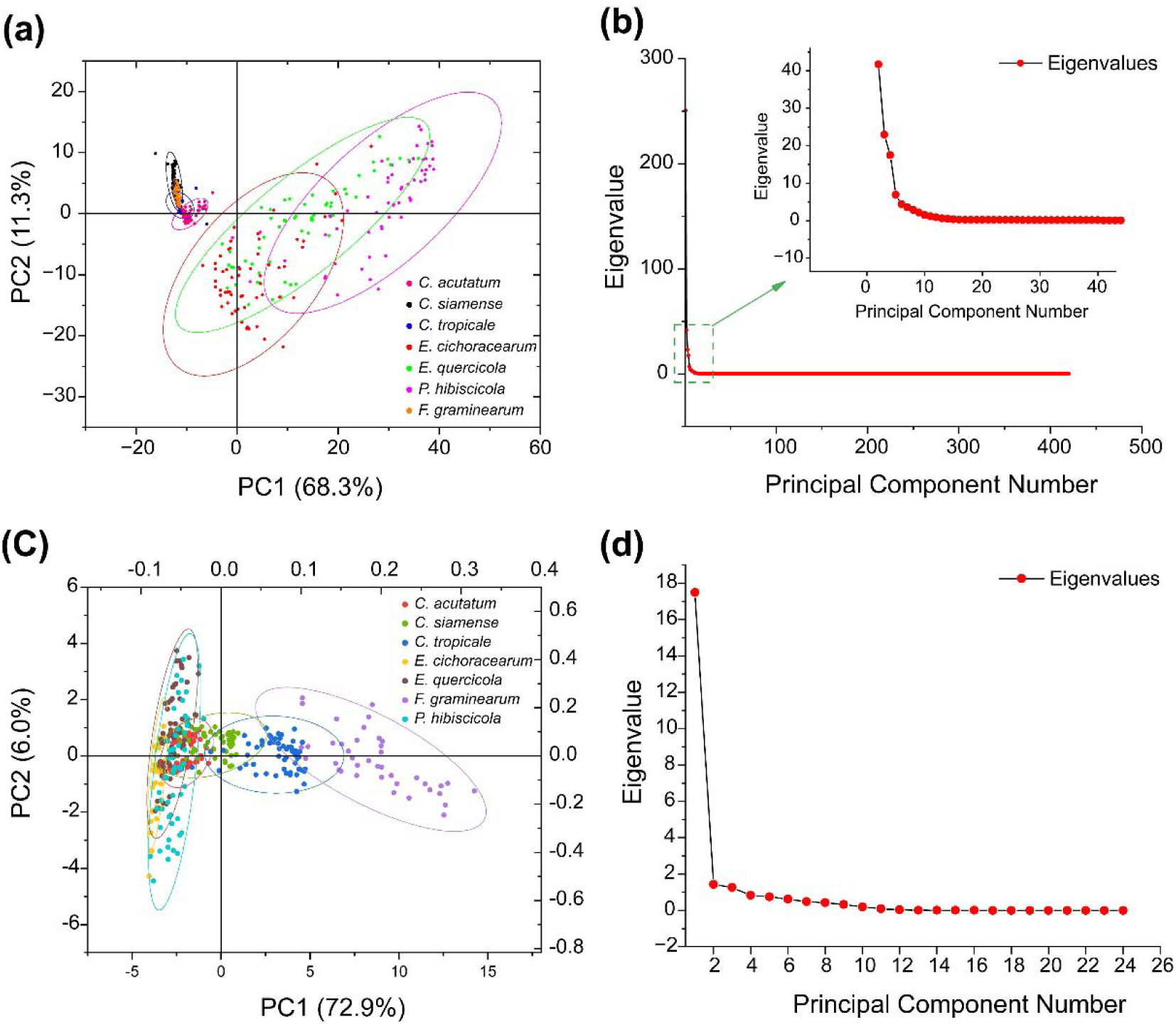
PCA on spectra of the seven processed species. (a) Score plot of the Raman spectra based on full spectral range. (b) Scree plot and partial enlargement corresponding to (a). (c) Score plot of the Raman spectra based on features generated from the eight characteristic wavenumber ranges. (d) Scree plot corresponding to (c). It should be noted that ellipses in (a) and (c) represent 95% confidence intervals.

Another PCA was conducted using the uniform features generated from the eight characteristic wavenumber ranges (Fig. 1). Compared with the PCA based on the whole spectral range, it achieved better classification for *F. graminearum*, with slight overlap with *C*. *tropicale* but clear separation from others. In the three *Colletotrichum* species clusters, a notable distinction between *C*. *acutatum* and *C*. *tropicale* was observed, but *C*. *siamense* became a significant interference. Furthermore, the three powdery mildew species clearly overlapped within the 95% confidence interval and were confounded with *C*. *acutatum* (Fig. 3c). As shown in the PCA scree plot (Fig. 3d), the eigenvalue at the inflection point is 2, indicating that the first two principal components have captured most of the features, and that more principal components would not provide more significant improvement in clustering.

In recent years, Raman spectroscopy has been applied to identify various fungal pathogens such as those related to fruit spoilage [24], Russulales [61], dermatophyte [22], and respiratory diseases [23]. As the most commonly used method to analyze Raman spectra of fungal conidia, PCA has demonstrated acceptable performance of clustering at the genus level [19,21,22,24,62,63]. However, as shown by the overlap in the PCA based on raw and processed Raman spectra, it remains challenging to accurately identify fungal at the species level. Thus, there is an urgent need for faster and more reliable identification of fungal pathogens from Raman spectral data that outperforms the existing PCA method.

### 3.3 Validation of data-driven models

SVMs, DTs, and XGBoost were trained for efficient identification using the features generated from the eight characteristic wavenumber ranges (Fig. 1). Six-fold cross-validation was performed to evaluate the model performance and optimize the model hyperparameters. Namely, for each species, 40 of the 48 samples in the training set were used to train a model, while the remaining 8 samples served as a validation set to evaluate the performance of the trained model. This process was repeated six times to select the models that achieved the most accurate classification.

The SVM with a kernel of linear function showed the best macro-average precision (MAP = 0.85) compared to those with a kernel of radial (MAP = 0.81), polynomial (MAP = 0.62), and Sigmoid (MAP = 0.42) functions (Fig. 4a). A linear kernel function allowed SVMs to generate a hyperplane to isolate the samples in the feature space with a certain number of dimensions (e.g., 24 dimensions for the feature space in this study) [28]. Thus, a 23-dimensional hyperplane achieved acceptable classification of conidial samples based on the transformed dataset. The other three kernel functions can increase the dimensions of the feature space (e.g., addition of a new feature that equals the sum or product of two original features). However, dimensionality expansion did not improve the classifiability of the dataset likely due to the overfitting caused by nonlinear kernels.

**Fig. 4.**
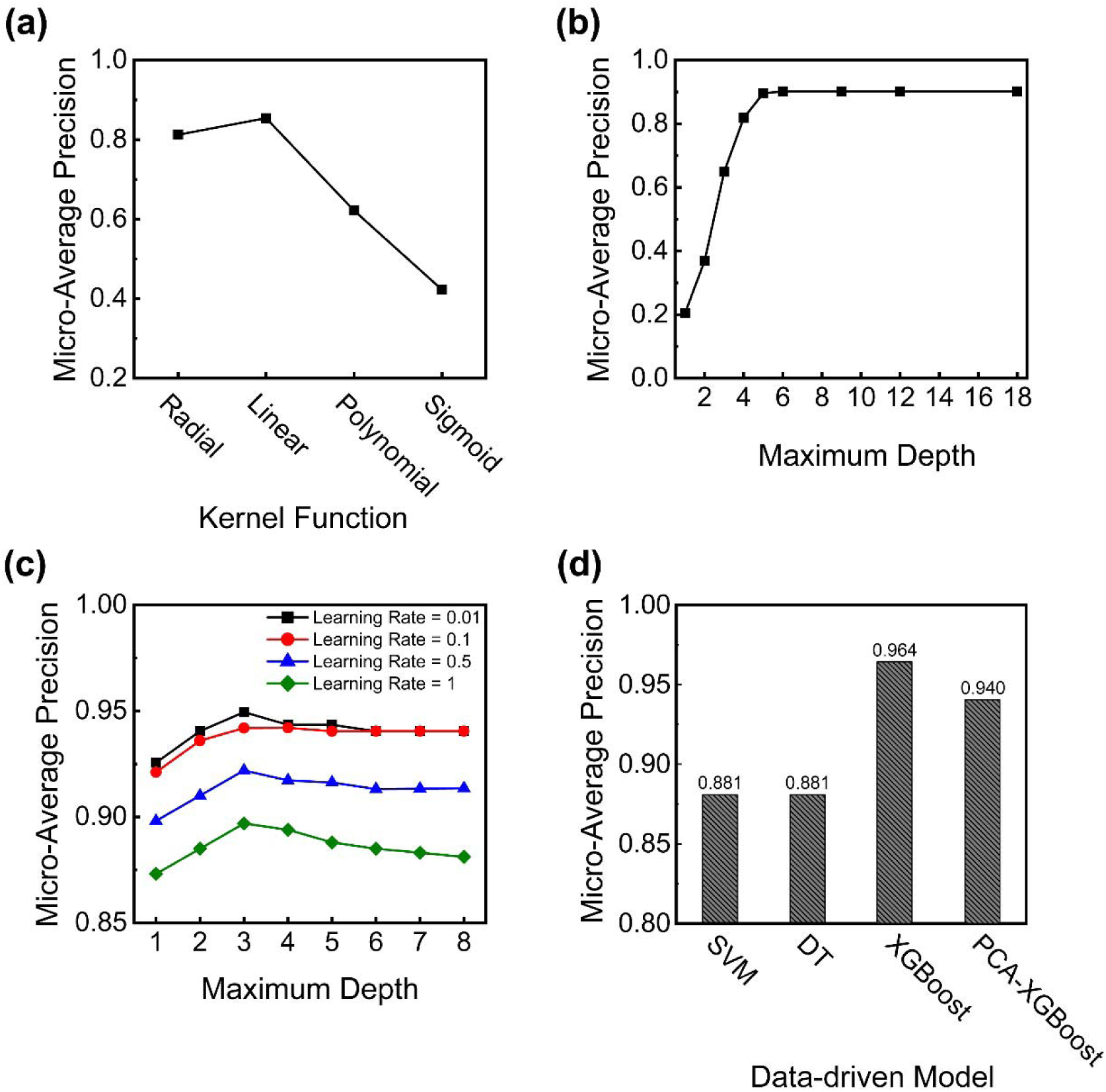
Training and optimization of (a) SVMs with different kernel functions, (b) DTs with different maximum depth, and (c) XGBoosts with maximum depth and learning rate. (d) is the comparison of testing results between the optimal SVM, DT, XGBoost, and the control PCA-XGBoost.

DTs can improve the prediction accuracy by increasing the depth as more decision steps (i.e., nodes) can facilitate the partitioning of the samples (Fig. 4b). The DT reached the optimal performance of MAP = 0.90 when the maximum depth was set to six, which was slightly higher than the optimal SVM. Additional decision steps did not further improve the classification. Moreover, a complex DT structure could lead to overfitting [64,65]. In line with Occam’s Razor Principle, the tree with a maximum depth of six was selected as the optimal DT, as it had the most sparse/pruned structure while achieving the highest prediction accuracy [28].

XGBoost as a forest-based algorithm introduces a regularization term of forest complexity into the loss function [32], which can efficiently mitigate model overfitting by controlling the structural complexity of DTs. Compared to SVMs and DTs, XGBoost achieved the highest MAP of 0.95 when the maximum depth and learning rate were set to 3 and 0.01, respectively (Fig. 4c). A low learning rate increased the precision but consumed more computational power. For example, the runtime of XGBoost increased from second and minute levels when the learning rate was lowered from 1 and 0.01. In practical applications, a lightweight model is preferred if it can provide comparably accurate predictions. Therefore, XBGoost with a maximum depth of 3 and a learning rate of 0.01 was selected as the optimal model.

To evaluate the effect of feature selection on classification, a PCA-XGBoost was also trained with a dataset composed of principal components derived from the PCA on full spectra. The PCA-XGBoost was constructed with different numbers of principal components, and those with more features tended to generate more accurate classification. Based on cross-validation, the model configured with a learning rate of 0.01, depth of 5, and inputs of 20 components yielded the best precision of 0.93 (Fig. S2).

### 3.4 Testing of data-driven models

The remaining 20% of the dataset (12 sample for each species, in total 84 samples for the seven species) was used to test the optimized data-driven models. SVM and DT achieved the same prediction precision (MAP = 0.88). Consistent with the cross-validation results, XGBoost generated the more accurate classification with MAP of 0.96 (Fig. 4d), while PCA-XGBoost achieved a comparable precision of 0.94. In both model validation and testing, the PCA-XGBoosts with inputs of 20 principal components generated less accurate classification than the XGBoosts (t-test, *p* < 0.05). This might result from the loss of information during data orthogonalization to the principal directions of the PCA, which increased the difficulty in learning the characteristics of the dataset [36,37]. Meanwhile, unlike the features directly generated from the raw Raman shifts, the principal components of PCA have lost their physicochemical properties and, consequently, cannot be used to interpret the correlations between features and labels.

All three data-driven models accurately classified *F. graminearum* and the three *Colletotrichum* species, with only two incorrect classifications of *C*. *siamense* (Supplementary File 3). On the other hand, powdery mildews were not consistently classified. In particular, the models often confused *P*. *hibiscicola* with *E*. *quercicola*, or *E*. *cichoracearum* with *E*. *quercicola* (Supplementary File 3). The inaccurate classification of powdery mildews might be attributed to the considerable intraspecies diversity (Fig. 2), which could result in highly uniform spectra from heterospecific individuals with indistinguishable morphologies (Fig. S3) and intracellular substances [66,67]. For example, all three data-driven models failed to classify a sample of *E*. *cichoracearum* (Sample #234), likely due to the high similarity between this sample and *E. quercicola* (Supplementary File 3). Nevertheless, successful classification of most of the samples highlighted the advantage of combining Raman spectroscopy and data-driven modeling over conventional classification methods such as those based on morphology and molecular biology. For example, multi-gene sequence analysis can achieve species-level identification[68,69], but the procedure is much more laborious, time-consuming, and expertise-intensive compared to Raman coupled with data-driven classification.

The results collectively show simpler and more reliable classification of fungal conidia by data-driven models than by PCA. Raman spectra can consist of noise and redundant signals that lead to overfitting and compromise predictive performance. Manual extraction of features from Raman shifts represents a promising approach to retain sufficient characteristics for accurate classification. On the other hand, PCA as a common approach for feature extraction and noise filtering did not improve data-driven classification of fungal conidia [44,45]. This likely resulted from the two inherent limitations of dimensionality reduction in PCA: 1) some of significant characteristics might be omitted during the process, and 2) the new features were generated through composite transformations from the raw datasets and subsequently lost interpretability [36,37]. Therefore, for the datasets with clear physical significance, over-transformation may not be beneficial. To further improve data-driven classification, additional features (e.g., features from other Raman spectral shifts) could be selected, and more samples, especially those with low prediction precision (e.g., *P. hibiscicola*), should be collected.

## 4. Conclusion

In this study, a Raman spectrum-based data-driven modeling approach was developed to identify phytopathogenic conidia. The first step of the approach was to characterize the Raman spectra, and the second step was to train data-driven models to classify the samples. In this study, eight fungal species were characterized using Raman, during which three common primary Raman peaks and several minor peaks were observed. Based on the features extracted from the spectral data, three data-driven models, SVMs, DTs, and XGBoost, were trained. SVMs and DTs achieved a prediction precision of 0.88, while XGBoost exhibited exceptional prediction performance with a precision of 0.96. The comparison between XGBoost and PCA-XGBoost indicated that features extracted from the raw datasets were more favorable in terms of model training than those extracted with PCA. This modeling approach represents a rapid, reliable, and accurate method to identify phytopathogenic conidia at the species level.

## Acknowledgment

This work was financially supported by Hainan Province Science and Technology Talent Innovation Project (KJRC2023B14); National Natural Science Foundation of China (32360640); Collaborative Innovation Center of Nanfan and High Efficiency Tropical Agriculture of Hainan University (XTCX2022NYA01); The U.S. Department of Agriculture (2020-67019-31027).

## Data Availability

The original Raman spectral data are available from MicrobioRaman database with accession number S-MBRS11 (https://www.ebi.ac.uk/biostudies/microbioraman/studies). Processed Raman spectral data we used in this manuscript has attached as Supplementary Data.

## References

[1] Chatterjee S, Kuang Y, Splivallo R, Chatterjee P, Karlovsky P. Interactions among filamentous fungi *Aspergillus niger*, *Fusarium verticillioides* and *Clonostachys rosea*: fungal biomass, diversity of secreted metabolites and fumonisin production. BMC Microbiol 2016; 16:83. 10.1186/s12866-016-0698-3.

[2] Fisher MC, Henk DanielA, Briggs CJ, Brownstein JS, Madoff LC, McCraw SL, et al. Emerging fungal threats to animal, plant and ecosystem health. Nature 2012; 484:186–94. 10.1038/nature10947.

[3] Fisher MC, Gurr SJ, Cuomo CA, Blehert DS, Jin H, Stukenbrock EH, et al. Threats posed by the fungal kingdom to humans, Wildlife, and Agriculture. MBio 2020; 11. 10.1128/mBio.00449-20.

[4] Doehlemann G, Ökmen B, Zhu W, Sharon A. Plant pathogenic fungi. Microbiol Spectr 2017; 5. 10.1128/microbiolspec.FUNK-0023-2016.

[5] Peng Y, Li SJ, Yan J, Tang Y, Cheng JP, Gao AJ, et al. Research progress on phytopathogenic fungi and their role as biocontrol agents. Front Microbiol 2021; 12. 10.3389/fmicb.2021.670135.

[6] Wang F, Sethiya P, Hu X, Guo S, Chen Y, Li A, et al. Transcription in fungal conidia before dormancy produces phenotypically variable conidia that maximize survival in different environments. Nat Microbiol 2021; 6:1066–81. 10.1038/s41564-021-00922-y.

[7] Nezhad AS. Future of portable devices for plant pathogen diagnosis. Lab Chip 2014; 14:2887–904. 10.1039/C4LC00487F.

[8] McCartney HA, Foster SJ, Fraaije BA, Ward E. Molecular diagnostics for fungal plant pathogens. Pest Manag Sci 2003; 59:129–42. 10.1002/ps.575.

[9] Pryce TM, Palladino S, Kay ID, Coombs GW. Rapid identification of fungi by sequencing the ITS1 and ITS2 regions using an automated capillary electrophoresis system. Med Mycol 2003; 41:369–81. 10.1080/13693780310001600435.

[10] Ko Ko TW, Stephenson SL, Bahkali AH, Hyde KD. From morphology to molecular biology: can we use sequence data to identify fungal endophytes? Fungal Divers 2011; 50:113–20. 10.1007/s13225-011-0130-0.

[11] Raja HA, Miller AN, Pearce CJ, Oberlies NH. Fungal identification using molecular tools: a primer for the natural products research community. J Nat Prod 2017; 80:756–70. 10.1021/acs.jnatprod.6b01085.

[12] Capote N, Mara A, Aguado A, Snchez-Torres P. Molecular tools for detection of plant pathogenic fungi and fungicide resistance. Plant Pathol, InTech; 2012. 10.5772/38011.

[13] Midorikawa GEO, Miller RNG, Bittencourt DM de C. Molecular identification and detection of foodborne and feedborne mycotoxigenic fungi. Molecular Techniques in Food Biology, Wiley; 2018, p. 385–407. 10.1002/9781119374633.ch17.

[14] De Gelder J, De Gussem K, Vandenabeele P, Moens L. Reference database of Raman spectra of biological molecules. Journal of Raman Spectroscopy 2007; 38:1133–47. 10.1002/jrs.1734.

[15] Ivleva NP, Wagner M, Horn H, Niessner R, Haisch C. In situ surface-enhanced Raman scattering analysis of biofilm. Anal Chem 2008; 80:8538–44. 10.1021/ac801426m.

[16] Madzharova F, Heiner Z, Gühlke M, Kneipp J. Surface-enhanced hyper-Raman spectra of adenine, guanine, cytosine, thymine, and uracil. The Journal of Physical Chemistry C 2016; 120:15415–23. 10.1021/acs.jpcc.6b02753.

[17] Kuo M-T, Lin C-C, Liu H-Y, Yang M-Y, Chang H-C. Differentiation between infectious and noninfectious ulcerative keratitis by Raman spectra of human teardrops: a pilot study. Investigative Opthalmology & Visual Science 2012; 53:1436. 10.1167/iovs.11-7923.

[18] Ghosal S, Macher JM, Ahmed K. Raman microspectroscopy-based identification of individual fungal spores as potential indicators of indoor contamination and moisture-related building damage. Environ Sci Technol 2012; 46:6088–95. 10.1021/es203782j.

[19] Huang WE, Griffiths RI, Thompson IP, Bailey MJ, Whiteley AS. Raman microscopic analysis of single microbial cells. Anal Chem 2004; 76:4452–8. 10.1021/ac049753k.

[20] Huang WE, Li M, Jarvis RM, Goodacre R, Banwart SA. Shining light on the microbial world, 2010, p. 153–86. 10.1016/S0065-2164(10)70005-8.

[21] Gan Q, Wang X, Wang Y, Xie Z, Tian Y, Lu Y. Culture free detection of crop pathogens at the single cell level by micro Raman spectroscopy. Advanced Science 2017;4. 10.1002/advs.201700127.

[22] Witkowska E, Jagielski T, Kamińska A. Genus- and species-level identification of dermatophyte fungi by surface-enhanced Raman spectroscopy. Spectrochim Acta A Mol Biomol Spectrosc 2018; 192:285–90. 10.1016/j.saa.2017.11.008.

[23] Žukovskaja O, Kloß S, Blango MG, Ryabchykov O, Kniemeyer O, Brakhage AA, et al. UV-Raman spectroscopic identification of fungal spores important for respiratory diseases. Anal Chem 2018; 90:8912–8. 10.1021/acs.analchem.8b01038.

[24] Guo Z, Wang M, Barimah AO, Chen Q, Li H, Shi J, et al. Label-free surface enhanced Raman scattering spectroscopy for discrimination and detection of dominant apple spoilage fungus. Int J Food Microbiol 2021; 338:108990. 10.1016/j.ijfoodmicro.2020.108990.

[25] Alvi M, Batstone D, Mbamba CK, Keymer P, French T, Ward A, et al. Deep learning in wastewater treatment: a critical review. Water Res 2023; 245:120518. 10.1016/j.watres.2023.120518.

[26] Kalantar B, Pradhan B, Naghibi SA, Motevalli A, Mansor S. Assessment of the effects of training data selection on the landslide susceptibility mapping: a comparison between support vector machine (SVM), logistic regression (LR) and artificial neural networks (ANN). Geomatics, Natural Hazards and Risk 2018; 9:49–69. 10.1080/19475705.2017.1407368.

[27] Christopher M. Bishop. Pattern recognition and machine learning. Springer; 2006.

[28] Zhou Z-H. Dimensionality reduction and metric learning. Mach Learn, Singapore: Springer Singapore; 2021, p. 241–64. 10.1007/978-981-15-1967-3_10.

[29] Khan S, Ullah R, Khan A, Wahab N, Bilal M, Ahmed M. Analysis of dengue infection based on Raman spectroscopy and support vector machine (SVM). Biomed Opt Express 2016; 7:2249–56. 10.1364/BOE.7.002249.

[30] Teh SK, Zheng W, Ho KY, Teh M, Yeoh KG, Huang Z. Diagnosis of gastric cancer using near-infrared Raman spectroscopy and classification and regression tree techniques. J Biomed Opt 2008; 13:034013. 10.1117/1.2939406.

[31] Chen Y, Su Y, Ou L, Zou C, Chen Z. Classification of nasopharyngeal cell lines (C666-1, CNE2, NP69) via Raman spectroscopy and decision tree. Vib Spectrosc 2015; 80:24–9. 10.1016/j.vibspec.2015.06.004.

[32] Chen T, Guestrin C. XGBoost: a scalable tree boosting system. Proceedings of the 22nd ACM SIGKDD International Conference on Knowledge Discovery and Data Mining, New York, NY, USA: ACM; 2016, p. 785–94. 10.1145/2939672.2939785.

[33] Ren X, Guo H, Li S, Wang S, Li J. A novel image classification method with CNN-XGBoost model. Digital Forensics and Watermarking, vol. 10431, Springer, Cham; 2017, p. 378–90. 10.1007/978-3-319-64185-0_28.

[34] Liew XY, Hameed N, Clos J. An investigation of XGBoost-based algorithm for breast cancer classification. Machine Learning with Applications 2021; 6:100154. 10.1016/j.mlwa.2021.100154.

[35] Qi Z. The text classification of theft crime based on TF-IDF and XGBoost model. 2020 IEEE International Conference on Artificial Intelligence and Computer Applications (ICAICA), IEEE; 2020, p. 1241–6. 10.1109/ICAICA50127.2020.9182555.

[36] Aziz R, Verma CK, Srivastava N. Dimension reduction methods for microarray data: a review. AIMS Bioeng 2017; 4:179–97. 10.3934/bioeng.2017.1.179.

[37] Palo HK, Sahoo S, Subudhi AK. Dimensionality reduction techniques: principles, benefits, and limitations. Data Analytics in Bioinformatics, Wiley; 2021, p. 77–107. 10.1002/9781119785620.ch4.

[38] Wang Y, Wang C, Rajaofera MJN, Zhu L, Xu X, Liu W, et al. WY195, a new inducible promoter from the rubber powdery mildew pathogen, can be used as an excellent tool for genetic engineering. Front Microbiol 2020; 11. 10.3389/fmicb.2020.610252.

[39] Huang Y-S, Karashima T, Yamamoto M, Hamaguchi H. Molecular-level investigation of the structure, transformation, and bioactivity of single living fission yeast cells by time- and space-resolved raman spectroscopy. Biochemistry 2005; 44:10009–19. 10.1021/bi050179w.

[40] Lu X, Ren W, Hu C, Liu C, Li Z. Plasmon-enhanced surface-enhanced Raman scattering mapping concentrated on a single bead for ultrasensitive and multiplexed immunoassay. Anal Chem 2020; 92:12387–93. 10.1021/acs.analchem.0c02125.

[41] Tang J-W, Liu Q-H, Yin X-C, Pan Y-C, Wen P-B, Liu X, et al. Comparative analysis of machine learning algorithms on surface enhanced Raman spectra of clinical *Staphylococcus* species. Front Microbiol 2021; 12. 10.3389/fmicb.2021.696921.

[42] McGill JA, Ogunnaike BA, Vlachos DG. Efficient gradient estimation using finite differencing and likelihood ratios for kinetic Monte Carlo simulations. J Comput Phys 2012; 231:7170–86. 10.1016/j.jcp.2012.06.037.

[43] Christopher M. Bishop. Neural networks for pattern recognition. Oxford university press; 1995.

[44] Villa-Manríquez JF, Castro-Ramos J, Gutiérrez-Delgado F, Lopéz-Pacheco MA, Villanueva-Luna AE. Raman spectroscopy and PCA-SVM as a non-invasive diagnostic tool to identify and classify qualitatively glycated hemoglobin levels *in vivo*. J Biophotonics 2017; 10:1074–9. 10.1002/jbio.201600169.

[45] Zhang L, Li C, Peng D, Yi X, He S, Liu F, et al. Raman spectroscopy and machine learning for the classification of breast cancers. Spectrochim Acta A Mol Biomol Spectrosc 2022; 264:120300. 10.1016/j.saa.2021.120300.

[46] Bro R, Kjeldahl K, Smilde AK, Kiers HAL. Cross-validation of component models: a critical look at current methods. Anal Bioanal Chem 2008; 390:1241–51. 10.1007/s00216-007-1790-1.

[47] Grandini M, Bagli E, Visani G. Metrics for multi-class classification: an overview. ArXiv 2020.

[48] Wang X, Sun M-J, Liu J-X, Deng Y-G, Mo Y-X, Tao Z-H. Analysis of astaxanthin in *Phaffia rhodozyma* using laser tweezers raman spectroscopy. Spectroscopy and Spectral Analysis 2012; 32:2433–7. 10.3964/j.issn.1000-0593(2012)09-2433-05

[49] Sharma SK, Nelson DR, Abdrabu R, Khraiwesh B, Jijakli K, Arnoux M, et al. An integrative Raman microscopy-based workflow for rapid in situ analysis of microalgal lipid bodies. Biotechnol Biofuels 2015; 8:164. 10.1186/s13068-015-0349-1.

[50] Wood BR, Heraud P, Stojkovic S, Morrison D, Beardall J, McNaughton D. A portable Raman acoustic levitation spectroscopic system for the identification and environmental monitoring of algal cells. Anal Chem 2005; 77:4955–61. 10.1021/ac050281z.

[51] Liu J, Huang Q. Screening of astaxanthin-hyperproducing *Haematococcus pluvialis* using fourier transform infrared (FT-IR) and Raman microspectroscopy. Appl Spectrosc 2016; 70:1639–48. 10.1177/0003702816645605.

[52] Noothalapati Venkata HN, Shigeto S. Stable isotope-labeled Raman imaging reveals dynamic proteome localization to lipid droplets in single fission yeast cells. Chem Biol 2012; 19:1373–80. 10.1016/j.chembiol.2012.08.020.

[53] Lin C-C, Yang Y-M, Liao P-H, Chen D-W, Lin H-P, Chang H-C. A filter-like AuNPs@MS SERS substrate for *Staphylococcus aureus* detection. Biosens Bioelectron 2014; 53:519–27. 10.1016/j.bios.2013.10.017.

[54] Avalos J, Pardo-Medina J, Parra-Rivero O, Ruger-Herreros M, Rodríguez-Ortiz R, Hornero-Méndez D, et al. Carotenoid biosynthesis in *Fusarium*. Journal of Fungi 2017; 3:39. 10.3390/jof3030039.

[55] Naz T, Ullah S, Nazir Y, Li S, Iqbal B, Liu Q, et al. Industrially important fungal carotenoids: advancements in biotechnological production and extraction. Journal of Fungi 2023; 9:578. 10.3390/jof9050578.

[56] Zhao X, Liu Z, He Y, Zhang W, Tong L. Study on early rice blast diagnosis based on unpre-processed Raman spectral data. Spectrochim Acta A Mol Biomol Spectrosc 2020; 234:118255. 10.1016/j.saa.2020.118255.

[57] Dou T, Sanchez L, Irigoyen S, Goff N, Niraula P, Mandadi K, et al. Biochemical origin of Raman-based diagnostics of Huanglongbing in grapefruit trees. Front Plant Sci 2021;12. 10.3389/fpls.2021.680991.

[58] Maquelin K, Kirschner C, Choo-Smith L-P, van den Braak N, Endtz HP, Naumann D, et al. Identification of medically relevant microorganisms by vibrational spectroscopy. J Microbiol Methods 2002; 51:255–71. 10.1016/S0167-7012(02)00127-6.

[59] Schuster KC, Reese I, Urlaub E, Gapes JR, Lendl B. Multidimensional information on the chemical composition of single bacterial cells by confocal Raman microspectroscopy. Anal Chem 2000; 72:5529–34. 10.1021/ac000718x.

[60] Kaczor A, Turnau K, Baranska M. In situ Raman imaging of astaxanthin in a single microalgal cell. Analyst 2011; 136:1109. 10.1039/c0an00553c.

[61] De Gussem K, Vandenabeele P, Verbeken A, Moens L. Raman spectroscopic study of *Lactarius* spores (Russulales, Fungi). Spectrochim Acta A Mol Biomol Spectrosc 2005; 61:2896–908. 10.1016/j.saa.2004.10.038.

[62] Xie C, Mace J, Dinno MA, Li YQ, Tang W, Newton RJ, et al. Identification of single bacterial cells in aqueous solution using confocal laser tweezers Raman spectroscopy. Anal Chem 2005; 77:4390–7. 10.1021/ac0504971.

[63] Dina NE, Gherman AMR, Chiş V, Sârbu C, Wieser A, Bauer D, et al. Characterization of clinically relevant fungi via SERS fingerprinting assisted by novel chemometric models. Anal Chem 2018; 90:2484–92. 10.1021/acs.analchem.7b03124.

[64] Bramer M. Principles of data mining. Springer; 2007.

[65] Bertsimas D, Dunn J. Optimal classification trees. Mach Learn 2017; 106:1039–82. 10.1007/s10994-017-5633-9.

[66] Notingher I. Raman spectroscopy cell-based biosensors. Sensors 2007; 7:1343–58. 10.3390/s7081343.

[67] Watanabe TM, Sasaki K, Fujita H. Recent advances in Raman spectral imaging in cell diagnosis and gene expression prediction. Genes (Basel) 2022; 13:2127. 10.3390/genes13112127.

[68] Arzanlou M, Bakhshi M, Karimi K, Torbati M. Multigene phylogeny reveals three new records of *Colletotrichum* spp. and several new host records for the mycobiota of Iran. J Plant Prot Res 2015; 55:198–211. 10.1515/jppr-2015-0027.

[69] Bragança CAD, Damm U, Baroncelli R, Massola Júnior NS, Crous PW. Species of the *Colletotrichum acutatum* complex associated with anthracnose diseases of fruit in Brazil. Fungal Biol 2016; 120:547–61. 10.1016/j.funbio.2016.01.011.

