## Supplementary material for "Direct Feature Identification from Raman Spectra and Precise Data-driven Classification of Phytopathogens at Single Conidium-Species Level": Fig. S

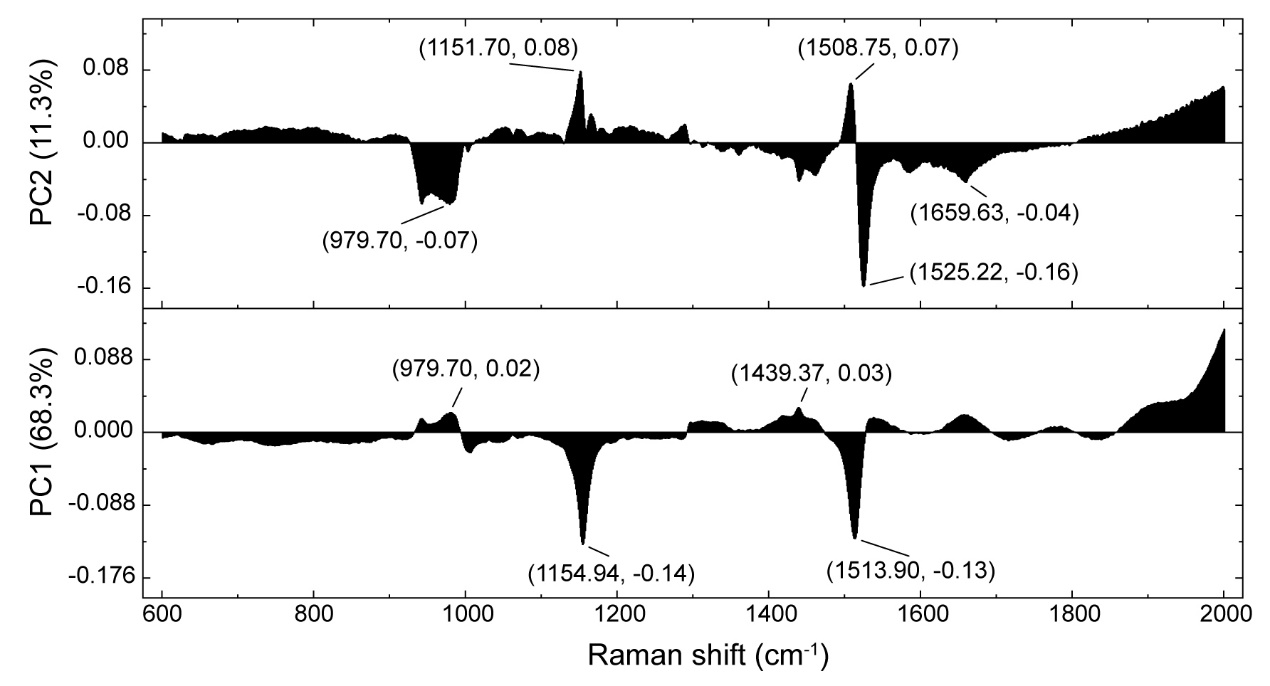

Fig. S1. Loading plots of PCA based on full spectral range. The black peaks and areas represent the basis for the discrimination of spectra.

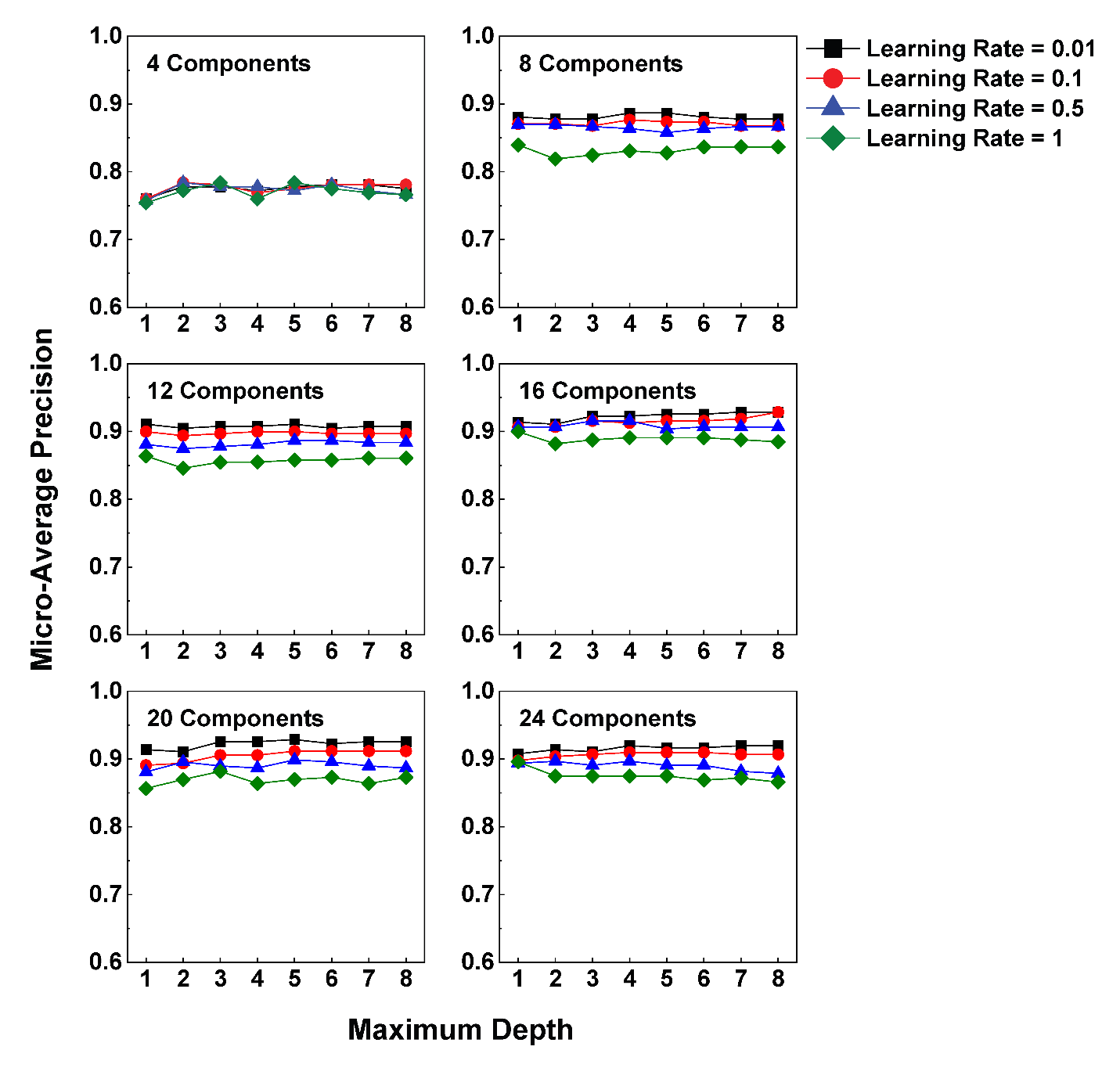

Fig. S2. The results from the hyperparameter tuning of PCA-XGBoost during cross-validation.

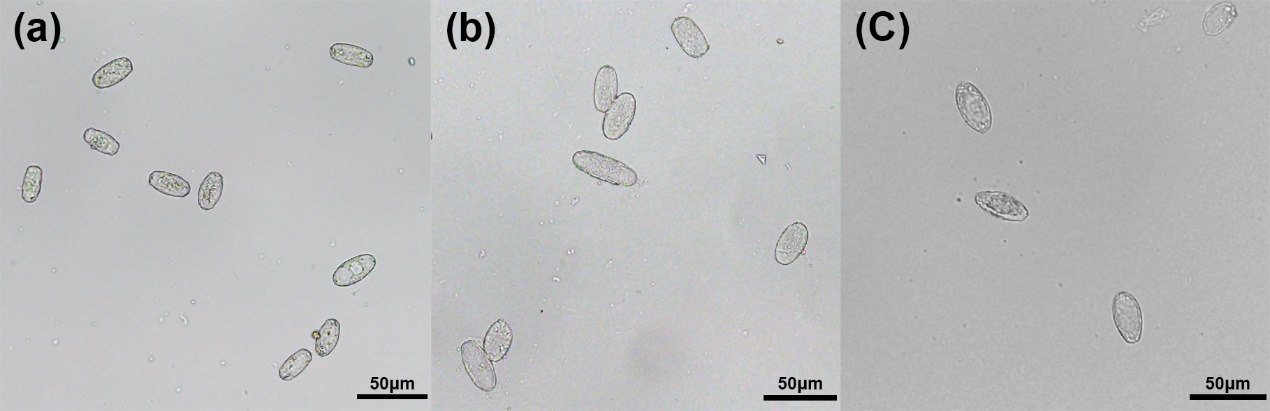

Fig. S3. The morphology of three powdery mildew conidia under microscopic observation. (a). *P. hibiscicola* (b) *E. cichoracearum*. (c) *E. quercicola*. The scale bar is 50 μm.
